# Translation Inhibition and Resource Balance in the Cell-Free Gene Expression System

**DOI:** 10.1101/142869

**Authors:** Vijayalakshmi H. Nagaraj, James M. Greene, Anirvan M. Sengupta, Eduardo D. Sontag

## Abstract

Quantifying the effect of vital resources on transcription and translation helps to understand the degree to which the concentration of each resource must be regulated for achieving homeostasis. Utilizing the synthetic transcription-translation (TX-TL) system, we study the impact of nucleotide triphosphates (NTPs) and magnesium (Mg^2+^), on gene expression. Recent observations of the counterintuitive phenomenon of suppression of gene expression at high NTP concentrations have led to the speculation that such suppression is due to the consumption of resources by transcription, hence leaving fewer resources for translation. In this work, we investigate an alternative hypothesis: direct suppression of the translation rate via stoichiometric mismatch in necessary reagents. We observe NTP-dependent suppression even in the early phase of gene expression, contradicting the resource limitation argument. To further decouple the contributions of transcription and translation, we performed gene expression experiments with purified mRNA. Simultaneously monitoring mRNA and protein abundances allowed us to extract a time-dependent translation rate. Measuring translation rates for different Mg^2+^ and NTP concentrations, we observe a complex resource dependence. We demonstrate that translation is the rate-limiting process that is directly inhibited by high NTP concentrations. Additional Mg^2+^ can partially reverse this inhibition. In several experiments, we observe two maxima of the translation rate viewed as a function of both Mg^2+^ and NTP concentration, which can be explained in terms of an NTP-independent effect on the ribosome complex and an NTP-Mg2^+^ titration effect. The non-trivial compensatory effects of abundance of different vital resources signals the presence of complex regulatory mechanisms to achieve optimal gene expression.

## 1 Introduction

The E. coli cell-free TX-TL system [1, 2, 3, 4, 5] is a promising tool for characterizing genetic networks in a minimally complex environment, and has been proposed as a technique for large scale industrial production of bioproducts ranging from synthetic fuels to drugs [6]. As a design tool, this technique allows biological engineers to explore genetic circuits in an analogous way to that in which electrical engineers use breadboards to understand electronic circuits. From a fundamental biology standpoint, the greater control provided over experimental conditions allows one to obtain a quantitative understanding of gene expression dynamics.

The experimental conditions in the TX-TL expression system must be precisely calibrated for optimal expression. Not only do various ion concentrations need to be accurately prescribed, but even resources like NTPs have to be optimized carefully for significant expression to occur. Our understanding of the resource requirements of gene expression is challenged by the observation that gene expression decreases with NTP concentration, once NTP is increased beyond a certain threshold [7]. This phenomenon is unexpected, since higher amounts of NTPs imply a greater source of energy available for transcription and translation. Siegal-Gaskin et al. [7] speculate that increased NTP levels enhance transcriptional activity, and thus exhaust most of the present NTPs. As a result, very little resources remain for translation to proceed, explaining the reduction of gene expression. In our investigations we contrast this hypothesis with a simpler explanation involving a direct suppression of gene expression, specifically that of the translation rate as a function of the overabundance of certain reagents.

In our system, which has no significant protein degradation mechanism, conversion of ATP to ADP, and the subsequent rise of ADP/ATP ratio [8], slow down protein production rates. A few hours into the experiment, this slowdown causes protein levels to saturate. Analyzing the gene expression profile thus requires us to be cognizant of the temporal dependence of transcription and, particularly, translation rates. We have developed methods of extracting such time-dependent rates.

One could reasonably believe that at times prior to saturation, resource limitation is peripheral to the underlying dynamics of gene expression. Therefore monitoring the protein and RNA levels at early times as a function of NTP and other reagents allows us to differentiate protein level suppression at high NTP concentrations. In general, we will use this experimental strategy to explore the landscape of resource dependence in the TX-TL system.

## 2 Materials and methods

### 2.1 Cell-free reaction conditions

Crude extract was prepared from BL21-Rosetta 2 strain, following the protocol of Sun et al. [5]. Basic Cell-Free reaction consists of a crude cell extract from E. coli (which contains endogenous transcription-translation machinery, mRNA and protein degradation enzymes plus all the soluble proteins), buffer and DNA. The amino acid mix was prepared following the protocol from [9]. All the experiments shown here are performed using the same batch of extract preparation to avoid variation between different batches. Reactions were carried out in a 10 μL volume using 1 to 2 nM plasmid, 10 to 20 μM malachite green to simultaneously monitor the RNA production. The reaction temperature was set to 29°C. Experiments were run for 14 to 20 hours with readings taken every 20 minutes, so that GFP and RNA expression dynamics could be monitored over a long period of time.

### 2.2 Monitoring RNA production

mRNA production was monitored using a plasmid provided by Dr. Murray’s lab which consists of a 35 base (MGapt) aptamer sequence incorporated in the 3’ UTR of the fluorescent protein reporter gene, 15 bases downstream of the stop codon and capable of binding the dye malachite green and produces fluorescence signal which can be measured at 650 nm wavelength. Fluorescence measurements were made in a total of 6 wells for each condition in a Biotek plate reader at 20 min intervals using excitation/emission wavelengths set at 610/650 nm (MGapt) and 485/525 nm (deGFP). Error bars indicate standard error over 6 replicates. The reported protein production at each time point represents a 20 minute time interval.

### 2.3 Plasmids and Bacterial Strains

Plasmids were a kind gift from Dr. Murray’s lab. BL21-Rosetta 2 strain for making crude extract was obtained from Novagen. Plasmid DNAs used for measuring protein and RNA production were prepared using Qiagen Plasmid Midi prep kits.

### 2.4 mRNA Preparation

First PCR was performed on pIVEX2.3 PT7-deGFP-MGapt plasmid using T7 promoter and T7 terminator primers obtained from IDT. The linear PCR was used as starting material for setting up transcription reactions using T7 RNA polymerase (Cellscript).

## 3 Results and discussion

We begin by examining the time-dependent gene expression profiles for increasing NTP concentrations. We use a plasmid described in Siegal-Gaskin et al., kindly provided to us by the Murray lab, with a built-in RNA aptamer (35-base MGapt sequence, which contains a binding pocket for malachite green (MG) dye and a fluorescent protein for accomplishing the simultaneous measurement of RNA and protein [7]. The result of a typical experiment is shown in Figure 1.

**Figure 1:**
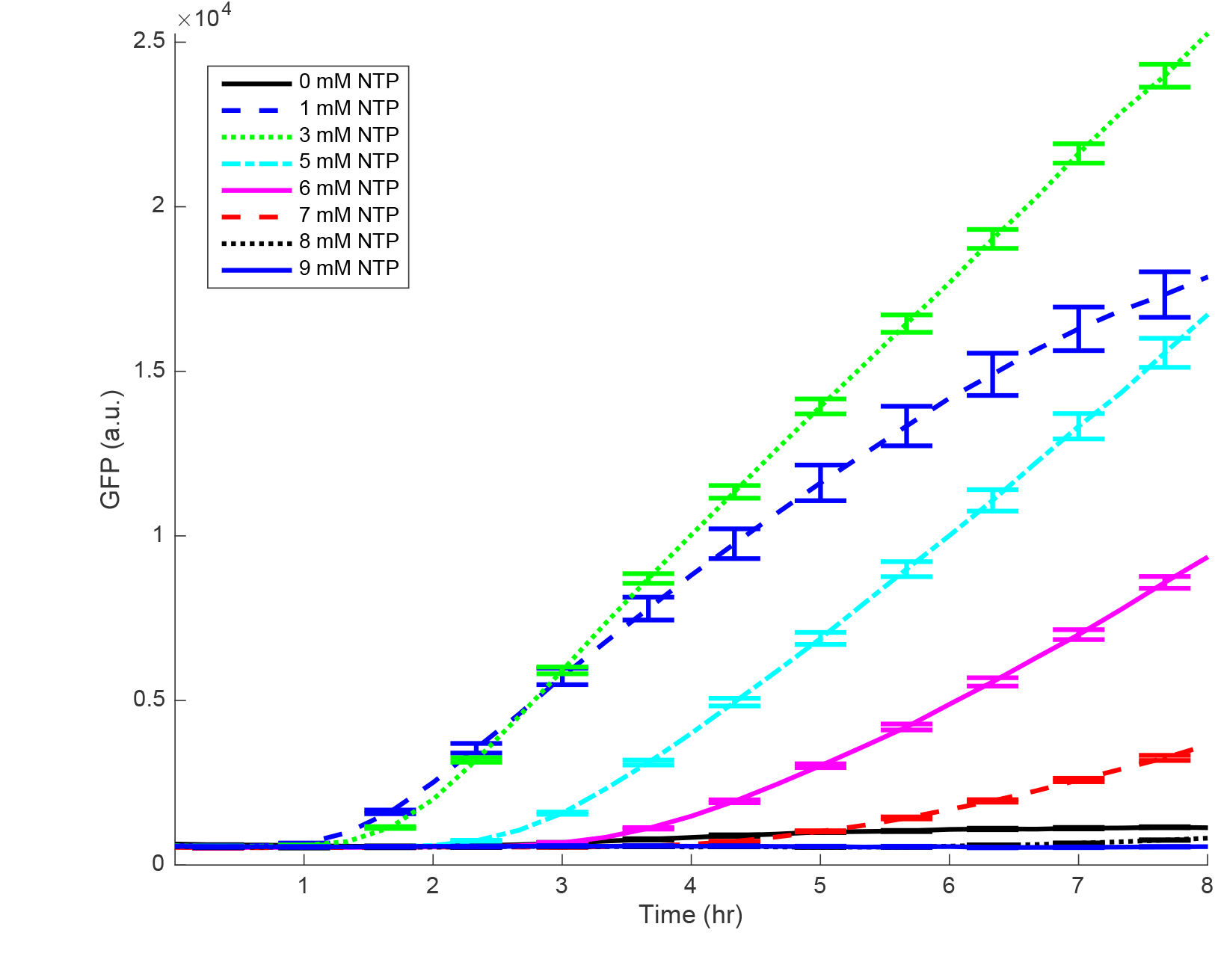
Translation kinetics demonstrating the GFP protein production for various NTP concentrations with DNA as the starting material. Data is measured every 20 minutes, while error bars, denoting one standard deviation, are plotted for every other data point for clarity. Mean and standard deviation are calculated over 6 replicates. Concentrations of Mg and K are 17.58 mM and 91.066 mM, respectively.

Figure 1 shows the time evolution of protein production as measured by GFP fluorescence, for various NTP concentrations. We illustrate only the initial 8 hours of the experiment, prior to saturation. Notice that protein production at early times is suppressed by increasing amounts of NTP. This observation conflicts with the hypothesis that resource limitation is driven by high transcriptional activity causing lower protein production. However, to further understand the roles of transcription and of translation, we proceeded to perform the experiments with the process of transcription eliminated. Below we describe these experiments.

We purified the mRNA from the plasmid mentioned above. Similarly to the transcription and translation process described previously, we measure the protein expression from purified mRNA added to the reaction mixture, in the absence of DNA. Therefore, in this experiment, the only processes affecting gene expression are mRNA decay and the time-dependent translation rate. The Mg binding 35 bp aptamer allows us to measure mRNA levels while GFP fluorescence gives us access to protein abundance. The resulting mRNA and protein profiles are shown in Figure 2 and Figure 3, respectively. RNA profiles in Figure 2 show approximate exponential decay, as expected.

**Figure 2:**
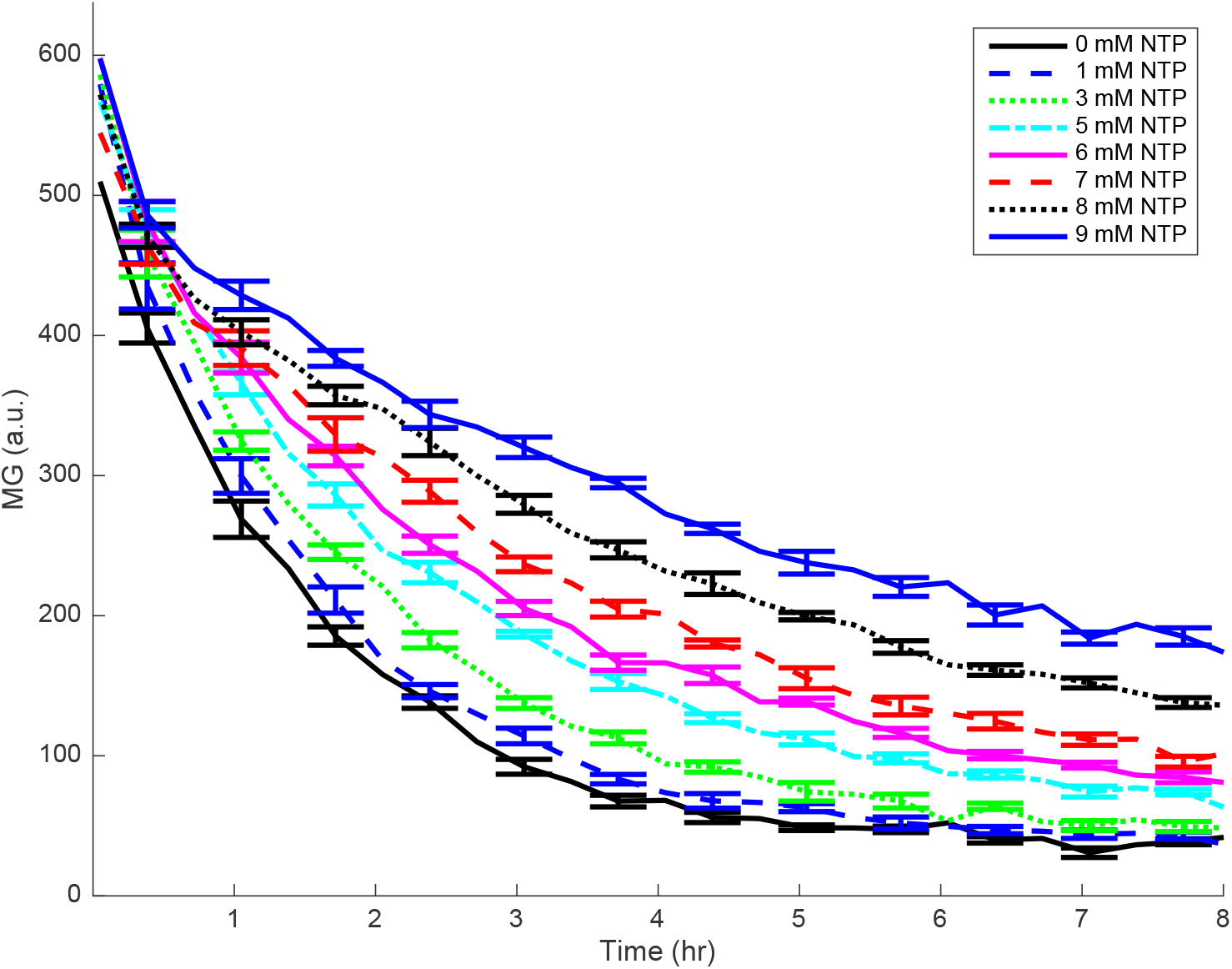
RNA decay profile for various NTP concentrations as a function of time, with mRNA as the starting material. Data is measured every 20 minutes, while error bars, denoting one standard deviation, are plotted for every other data point. Mean and standard deviation are calculated over 6 replicates. Concentrations of Mg and K are 17.58 mM and 91.066 mM, respectively.

**Figure 3:**
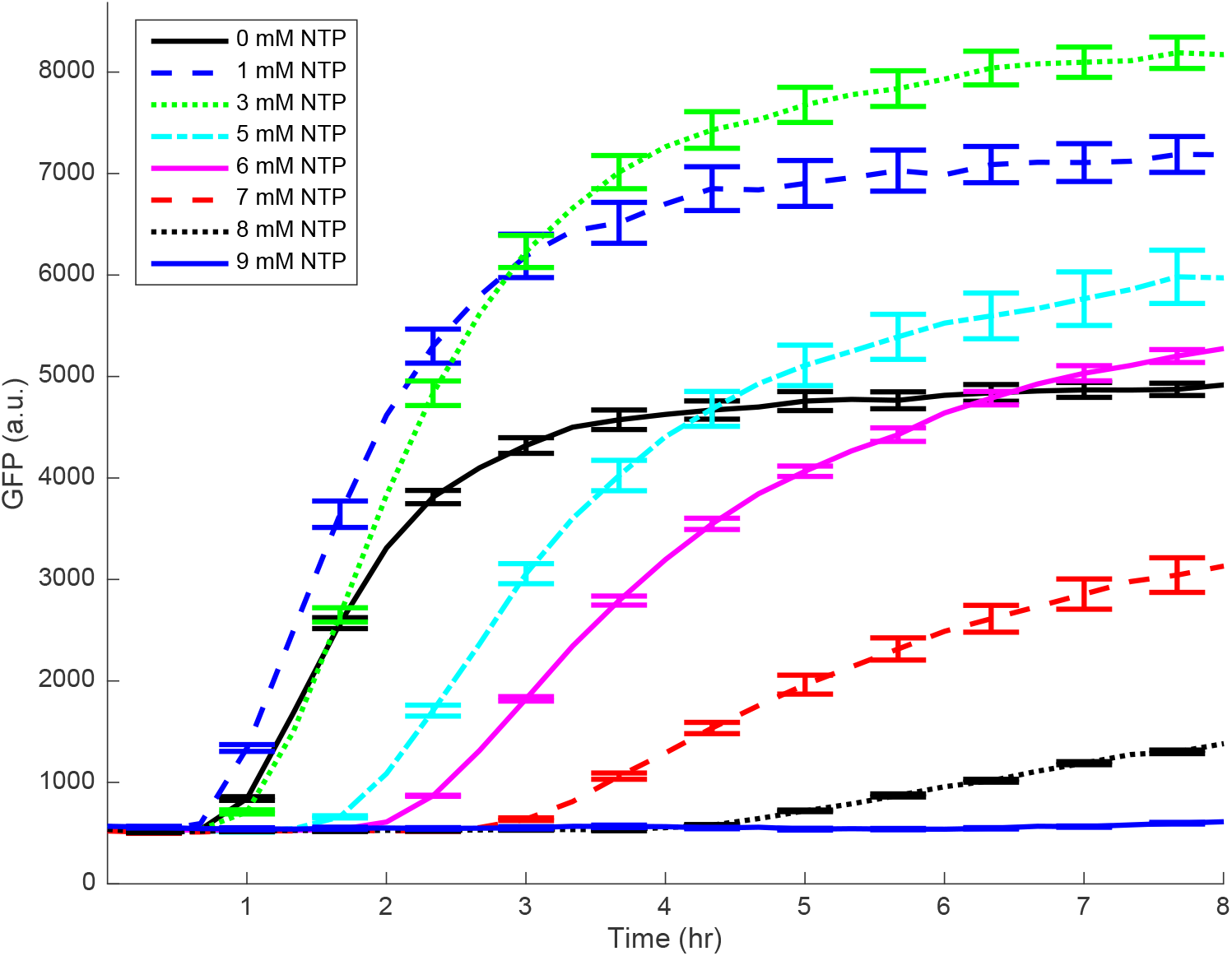
Translation kinetics showing the GFP protein production for various NTP concentrations with mRNA as the starting material. Data is measured every 20 minutes, while error bars, denoting one standard deviation, are plotted for every other data point. Mean and standard deviation are calculated over 6 replicates. Concentrations of Mg and K are 17.58 mM and 91.066 mM, respectively.

Using both the RNA and protein profiles, we extract time-dependent translation rates, following methods similar to those appearing in Siegal-Gaskin et al. from 2014 [7]. We assume that GFP (*g*) and MG (*m*) are related to protein (*p*) and RNA (*r*) expression as follows:

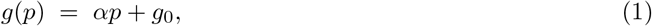

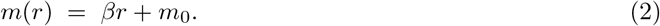

Here *g*_0_,*m*_0_ are parameters accounting for autofluorescence, and in general vary between experiments, while *α, β* are proportionality constants which convert between fluorescence measurements and the corresponding protein or mRNA concentrations. Importantly, we assume that *α* and *β* are fixed constants, independent of the experiment under consideration. Note that since RNA is unstable and undergoing approximate exponential decay (see Figure 2), we can estimate *m_0_* as the final temporal measurement of *m* for each experiment.

The fundamental quantity of interest is the (generally) time-varying translation rate *k_p_*(*t*). Since only translation is occurring, we assume that the rate of protein production is directly proportional to the amount of RNA *r*(*t*) at time t:

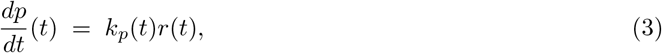

To estimate *k_p_*(*t*), we must first relate it to the observed quantities *g* and *m*. Defining

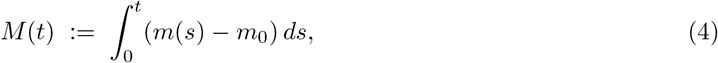

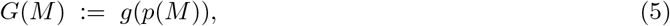

where *p*(*M*) is the protein concentration as a function of *M*, one can show (for details, including justification that *p* = *p*(*M*) is well-defined, see the Appendix) that *G*(*M*) satisfies the following ordinary differential equation (ODE):

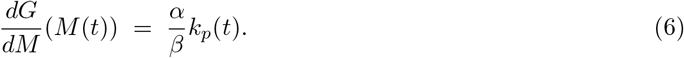

Since *α* and *β* are constant across experiments, equation (5) implies that 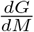 can be used as a proxy for the translation rate *k_p_*. Thus, to analyze the qualitative effects of Mg2^+^ and NTP on translation, we compute slopes in the G-M plane. See Figure 4 for a sample calculation, which we describe in the following.

**Figure 4:**
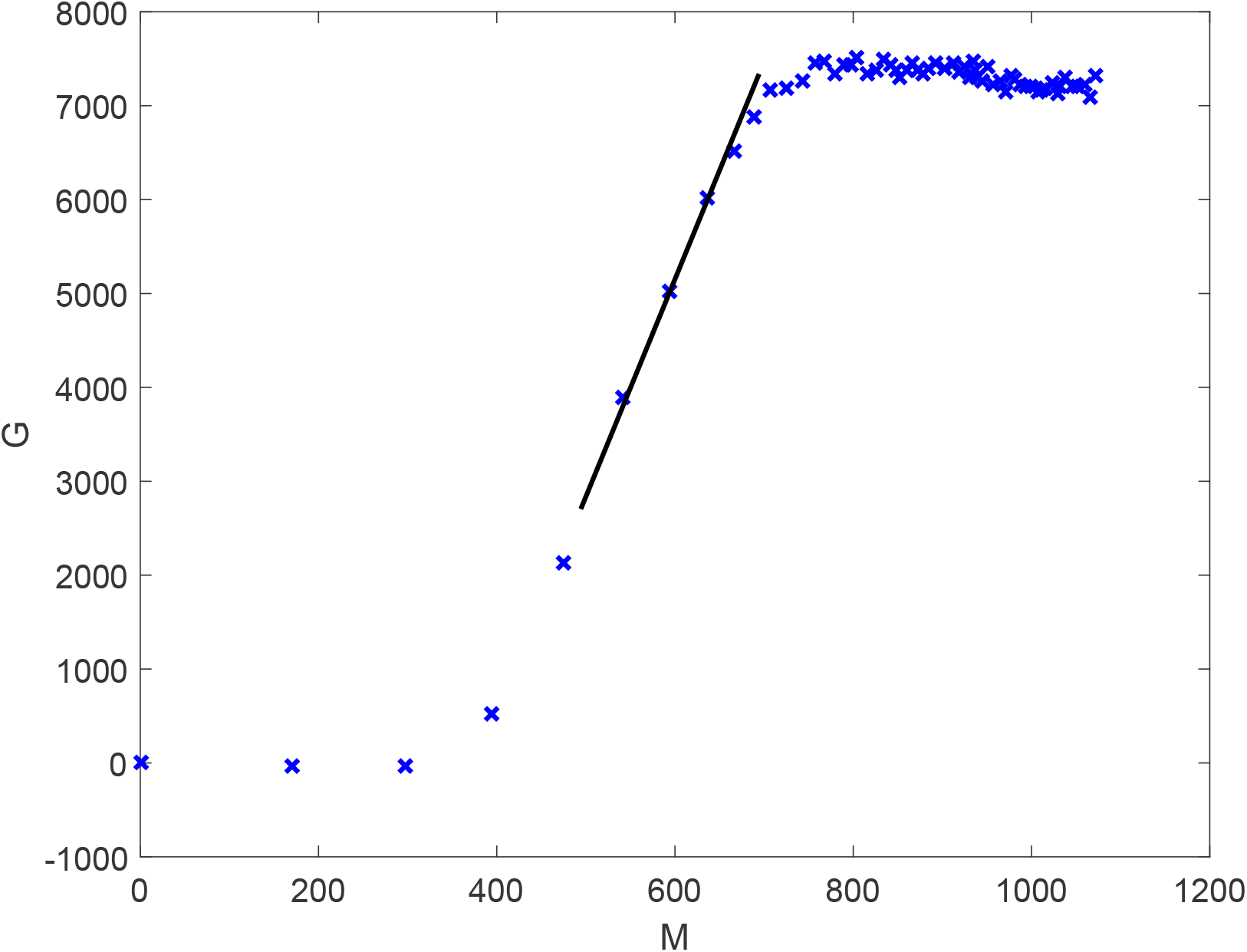
The change in protein vs. the integral of the RNA profile at 20.58 mM Mg2^+^ and no additional extra NTP. For most replicates and Mg/NTP concentrations, we observe a clear region of nearly constant translation rate. The extracted rate (i.e. slope) for this experiment is shown via the black solid line.

To calculate the translation rate (technically 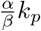) for each experiment, we observe the following general phenomenon. For all replicates, we observe a lag time *t*_0_ (see for example Figure 1 or 3) where protein levels appear to be approximately constant. Assuming this is generated by autofluorescence and thus is not due to protein production, translation has not yet begun to occur. This is verified in the *G – M* plane as in Figure 3, where a slope of approximately zero is observed initially. Afterwards, there exists a regime where translation occurs at a constant rate, which can be taken as proportional to *k_p_*(*t*) via equation (6). Towards the end of each experiment, as RNA is exhausted, protein production saturates, and again the slope of *G*(*M*) approaches zero. The initial translation rate is thus taken, in this work, as the slope of *G*(*M*) in the region of approximate linear growth. This behavior in the early and late regimes, taken together, implies that the slope should be largest during the period of linear growth, although some exceptions exist due to noise and the difficulty in computing derivatives with discrete data. To that end, we exclude early and late data from translation rate extraction, where these effects are most noticeable. The basis for our algorithm for computing *k*_p_ is then the following: transform to the (*M*, *G*) plane, compute moving windows of fixed length (5 data points were used, but results are robust to this value), use a least-squares regression to fit a line to the data, and measure the translation rate *k_p_*, as the maximum value over all such moving windows.

To verify that translation occurs, at least on an intermediate time window, at an approximate constant rate, we plot the GFP-MG (i.e. protein-RNA) translation data in the (*G*, *M*) plane for each experiment (each NTP and Mg2^+^ is performed with 6 replicates). For example, see the plot in Figure 4. Blue crosses represent experimental data points. In each case, we observe rapid linear growth between regions of slower growth; the initial slow growth corresponds to the experimental delay, which implies approximately no change in protein production: 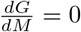, while the latter to the diminished RNA concentration in the well (i.e. saturation). This approximately linear regime is where we measure the translation rate *k_p_* via a least-squares regression (slope of black line). Note that the clustering of data points for large M values is due to the fact that the transformation *t* → *M*(*t*) is not an isometry. As time t increases, the decay of RNA (Figure 2) implies that the total amount of RNA (measured via M) stabilizes.

The computed translation rate is plotted against both NTP and Mg^2+^ concentrations in Figure 5, confirming our hypothesis that additional NTP is directly inhibiting translation.

**Figure 5:**
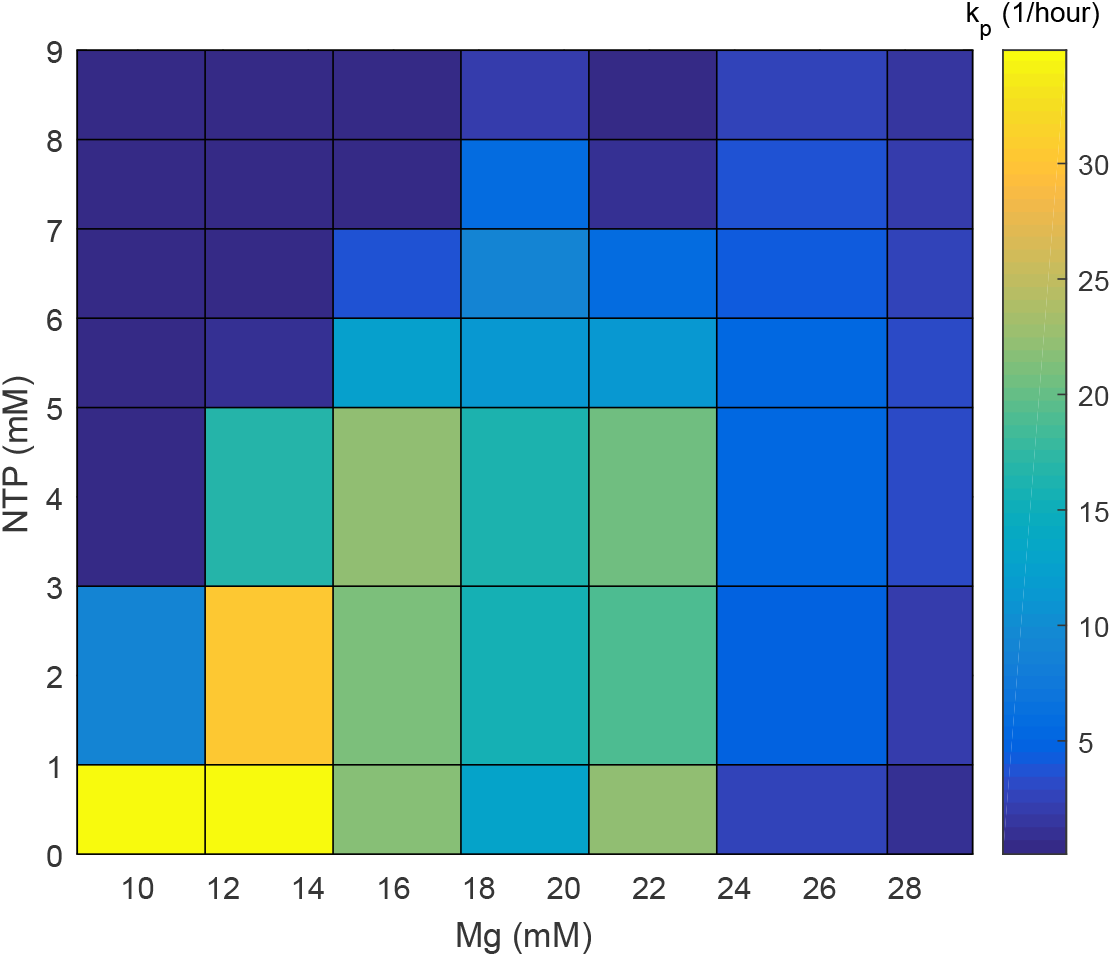
Surface plot showing the translation rate as a function of various NTP and Mg^2+^ concentrations.

Analyzing Figure 5, we observe a complex interdependence between Mg^2+^ and NTP concentrations on the translation rate dynamics. Specifically, for each value of NTP, two distinct maxima exist as a function of Mg^2+^. As the NTP concentration increases, one of these maxima shifts to increasing values of Mg^2+^. Recalling that Mg2^+^-NTP is the biologically relevant complex, we have developed a phenomenological model to explain both phenomenon: the shifting of a maximum, as well as the presence of multiple (here two) local maxima. Each phenomenon is described separately. We begin with a discussion of the shift of a (single) maximum. For the discussion of multiple maxima, see below.

In this model, a greater amount of NTP requires a greater amount of Mg^2+^ to be functional. Otherwise, the free NTP, unbound to Mg^2+^, ‘poisons’ the translation system. Initially, this may appear unintuitive in our experimental system, since the total concentration of NTP (a few mMs) is 2-3 times smaller than the total Mg^2+^ concentrations, with dissociation constant on the order of 0.01mM [10]. However, we recall that translation is affected not by the total ion concentration but instead by the free ions in the solution. Nierhaus et al. have shown [11] the binding of ribosomes to a significant fraction of Mg^2+^ and Potassium K, thereby leaving very little cytosolic ions available. The concentration of Mg available for binding with NTP to form NTP-Mg complex, which is central to gene expression [12], is therefore treated as an effective total concentration in our mathematical model.

More precisely, we assume that the translation rate takes the following general form:

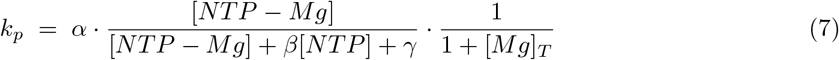

Here [*NTP – Mg*] denotes the concentration of the NTP-Mg complex, and is the driver of translation. The above expression represents the competition at the binding site between the unbound NTP and NTP-Mg molecules, with *β* and *γ* denoting (relative) dissociation constants, and *α* the maximum possible translation rate. We also have included a total Mg dependence of the form 1/(1 + [*Mg*]_*T*_), since it is known that Mg^2+^ in abundance will inhibit translation [13, 14, 15].

To calculate the concentrations used in the above formula, we assume all reactions are in equilibrium, and thus concentrations can be calculated via a steady-state analysis. We have the following association-dissociation reaction between Mg and NTP:

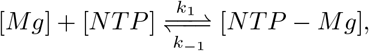

from which we can calculate the concentration of [*NTP – Mg*] as a function of [*Mg*]_*T*_, [*NTP*]_*T*_, and the dissociation constant *k_d_*(= *k*_−1_/*k*_1_). In general, we can show that for each total concentration of NTP, the above will possess a unique relative maximum, which increases as the total amount of NTP is increased. A sample plot is included in Figure 6 for demonstrative purposes.

**Figure 6:**
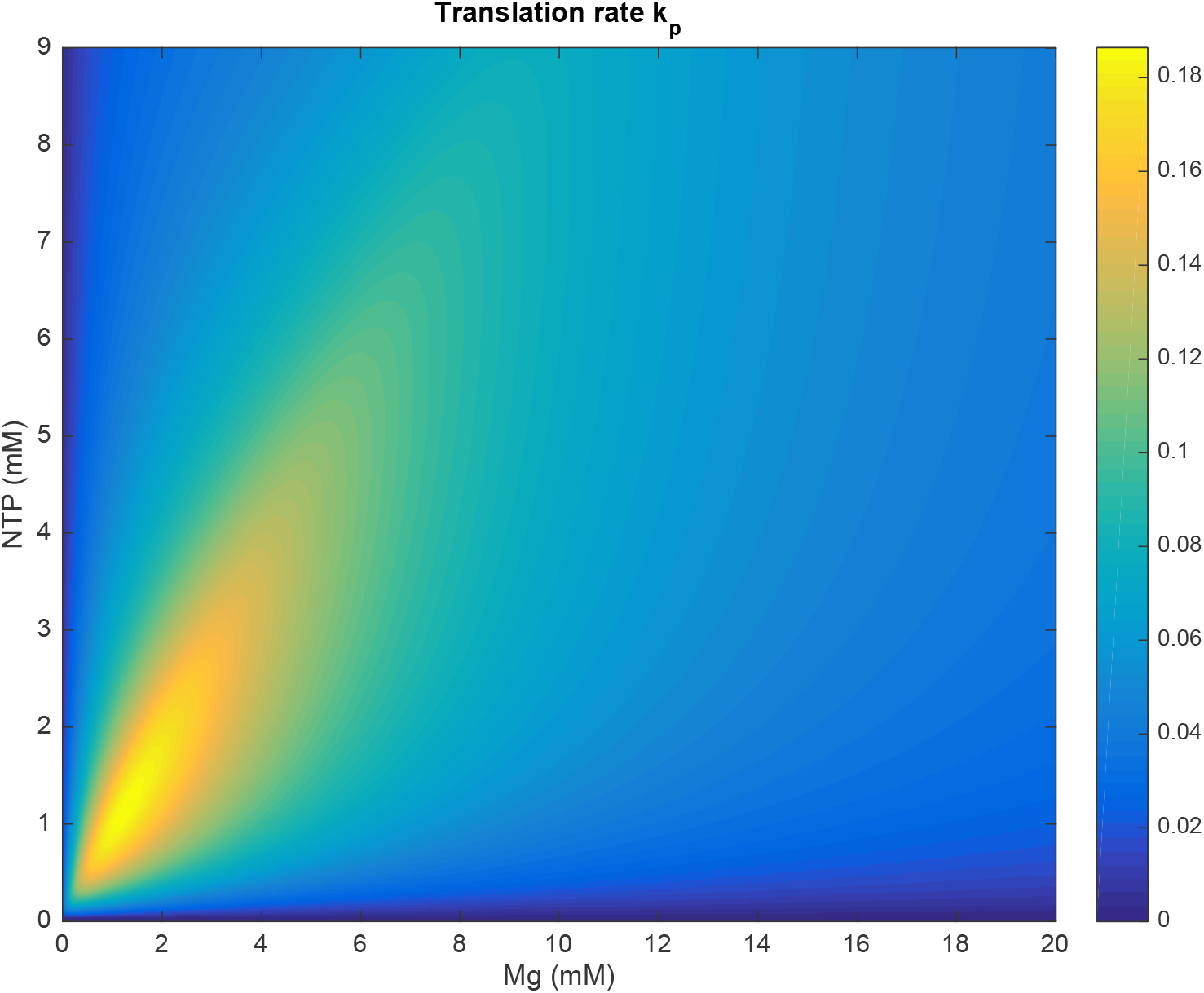
Heat map showing the translation rate as a function of both NTP and Mg^2+^ for the one maximum model (7).

The previous model does not capture the two-peak structure of the datasets. We believe that there are two different kinds of influence of Mg on translation. One of these has to do with NTP titration, as described before. The other has to do with stabilizing the structures of RNA and other molecules involved in the process [16, 11], and that effect has an optimal concentration requirement. The superposition of two such effects would then lead to two local maxima. Disentangling these two effects from the experimental data is indeed an interesting question.

Basic cellular processes like transcription and translation require many resources. As we try to build synthetic gene regulation networks, we need to understand competition for resources giving rise to limitations and tradeoffs. Both mathematical modeling and experimental studies are directed towards elucidation of this issue [17, 18, 7]. In the system studied, we provide evidence that suppression of gene expression due to added NTP is not primarily due to competition between transcription and translation as has been proposed in [7], but we suggest that the limitation of available free Mg^2+^ plays a role in this suppression. This suggestion is based on the observation that the suppression due to additional NTP is partly relieved by adding more Mg^2+^. In general, because of the nontrivial interaction between different resources, more studies need be done scanning multiple parameters simultaneously, while monitoring a particular phenotype.

Lastly, the fine-tuning needed in the TX-TL raises questions regarding the optimization of gene expression in vivo. Our knowledge of sensing and controlling levels of crucial reagents in the cell is currently incomplete at best, although quite a bit is known about homeostasis of ATP levels [19]. These synthetic biology studies therefore provide an important impetus for furthering our understanding.

## Author Contributions

Conceived of the project (E.D.S.), designed the experiments and analysis (E.D.S., A.M.S, and V.H.N.), performed the experiments and did preliminary data exploration (V.H.N.), analyzed data, especially, translation rate extraction (J.M.G.), performed mathematical modeling (J.M.G) and wrote the manuscript (V.H.N., J.M.G., A.M.S and E.D.S.).

## Acknowledgements

The authors thank Vincent Noireaux and members of the Murray lab for useful discussions and materials. This research is funded in part by grant ONR N00014-13-1-0074.

## Data availability

All raw data are available with the online version of this article.

## Supplementary data

Supplementary data are available at Synthetic Biology online.

## Appendix

In this section, we formally derive equation (6). Assume that the RNA concentration r(t) > 0 for all t in each experiment, and that translation can be described by (3):

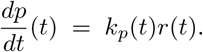

Equations (2) and (4) imply that

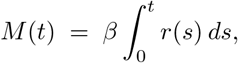

so that *M’*(*t*) = *r*(*t*) > 0,i.e. *M* is an increasing function of *t*. Thus, *M*(*t*) is invertible, and we can solve

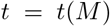

on some interval *M* ∈ [0, *M*_max_). Note that *M_max_* < ∞ as *r*(*t*) is exponentially decreasing, although this is not strictly necessary. Furthermore, we can compute the derivative of this function via the Inverse Function Theorem:

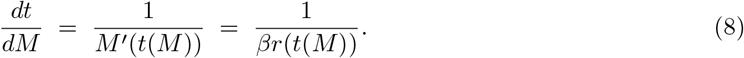

Defining the GFP marker as a function *M*,

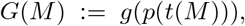

is thus well-defined, and we can compute its derivative using the chain-rule:

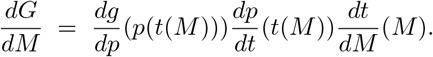

Equation (1) implies that 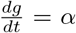, equation (3) that 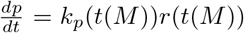, and equation (8) provides the last term on the right-hand side. Canceling *r*(*t*(*M*)) yields the expression

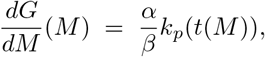

which is precisely (6), as desired.

